# Left mid-ventral temporal cortex interacts with early visual cortex and the anterior temporal lobe to support word individuation

**DOI:** 10.1101/411579

**Authors:** Matthew J. Boring, Elizabeth A. Hirshorn, Yuanning Li, Michael J. Ward, R. Mark Richardson, Julie A. Fiez, Avniel Singh Ghuman

## Abstract

The left mid-ventral temporal cortex (lmVTC) plays a dynamic role in reading. In this study we investigated the neural interactions that influence lmVTC dynamics and the lexical information these interactions are dependent on. We monitored activity with either intracranial electroencephalography or magnetoencephalography while participants viewed real words, pseudowords, consonant strings, and false fonts. A coarse level representation in early lmVTC activity allowed for decoding of visually dissimilar real words, pseudowords, and false fonts. Functional interactions between anterior ventral temporal regions, possibly containing stored knowledge about words, and low-order visual regions occurred after this initial stage of processing and was followed by the individuation of orthographically similar real words in lmVTC, but not similar pseudowords, letter strings, or false fonts. These results suggest that the individuation of real word representations in lmVTC is catalyzed by stored knowledge about word forms that emerges from network-level interactions with anterior regions of the temporal lobe.

## Introduction

A region in the left mid-ventral temporal cortex (lmVTC), sometimes referred to as the “Visual Word Form Area (VWFA),” responds preferentially to words over other object categories^1,2^ and is thought to play a key role in reading^1–6^. It has been shown that damage to^4,7–9^ or stimulation of^7,10^ the lmVTC can cause pure alexia, and there is evidence that reading expertise shapes response properties of the lmVTC, including differential activation to real words versus pseudowords (pronounceable but meaningless letter strings)^11–17^. However, there is still debate over the nature of orthographic representation in the lmVTC. Specifically, does the lmVTC encode whole-words^16,18,19^, sublexical features^3,7,20–23^, or purely visual statistics that are preferentially fed into higher-order language centers^5,24^?

A recent intracranial electroencephalography (iEEG) study demonstrated that early activity in the lmVTC only allowed the decoding of words that were orthographically dissimilar (hint vs. dome). The activity then evolved in a way that also allowed orthographically similar real words (hint vs. lint) to be disambiguated after 200 ms^7^. These results suggest that representations in the lmVTC are initially coarse but evolve over time, eventually allowing for the disambiguation of orthographically similar word forms. However, the degree to which this process is specific to known printed words, which have learned semantic and phonological associations in addition to their visual properties, has not yet been determined. Further, it is unknown if word individuation is achieved solely through hierarchical visual processing^18^ or is instead driven by interactions between the lmVTC and other parts of the language network that underpin phonological and/or semantic knowledge about words^17,25–27^.

The current study probes these questions by examining the lmVTC response to visually similar real words, pseudowords (pronounceable but meaningless letter strings), consonant strings (meaningless and unpronounceable letter strings), and false-fonts (orthographic stimuli of an unfamiliar alphabet) using source-localized magnetoencephalography (MEG) and iEEG. We hypothesized that if the disambiguation of orthographic representations in the lmVTC relies on learned semantic or phonological associations inherent to real words, then orthographically similar pseudowords and/or consonant strings would not be disambiguated by lmVTC activity. In contrast, if the refinement of lmVTC representation is independent of learned semantic or phonological knowledge, then we would expect to see a similar disambiguation for orthographically similar consonant strings and pseudowords. Additionally, we examined the functional connectivity between lmVTC and the rest of the cortex during the transition between coarse and individuated representations to assess the extent to which the disambiguation of orthographic representations is a network-level process.

Our results demonstrate that lmVTC responses to real words, pseudowords, consonant strings and false fonts allowed for reliable decoding of orthographically dissimilar stimuli from one another. However, while orthographically similar real words could be discriminated from lmVTC activity from 200-350 ms, orthographically similar pseudowords, consonant strings, and false fonts could not. Functional connectivity analysis in MEG revealed increased phase-locking between the lmVTC and left anterior temporal lobe and early visual cortex during the transition from these coarse to individuated real-word representations. Taken together, these results support the idea of early, coarse lmVTC representations that subsequently evolve through interactions with visual and semantic networks to allow for the disambiguation of orthographically similar real-words.

## Methods

### MEG data collection and preprocessing

#### Participants

16 participants gave written informed consent to participate in the MEG portion of the experiment consistent with protocol approved by the University of Pittsburgh’s Internal Review Board. One participant was removed from the analysis due to poor cortical surface reconstruction leaving 15 (5 males, ages 19-29) for the remaining analyses.

#### Experimental Paradigm

First, a category localizer consisting of words, hammers, houses and false-fonts was administered to identify word-selective cortical sources and constrain the word-individuation analysis. Then, a word individuation task was administered to probe the dynamics of word representation across different types of orthographic stimuli. For both the category localizer and word-individuation tasks stimuli were presented via custom scripted code in Psychtoolbox^28^ on a screen one meter in front of the participants. Stimuli occupied approximately 6 x 6° of visual angle and were shown for 300 ms with a variable inter-stimulus interval of approximately 1.5 s. One-sixth of the time a stimulus would be repeated, to which the participant responded with a button press. These trials were removed from the subsequent analyses. Three blocks of 140 trials each were performed for the category localizer and 5 blocks of 264 trials each were performed for the word individuation task. In total there were 90 trials per stimulus category in the category localizer and 30 trials per stimulus in the word-individuation task, after removing repeated trials.

In the word-individuation task, word and word-like stimuli consisted of four pairs of real words, pseudowords, false-fonts (Old Hungarian alphabet) and consonant-strings each differing from each other in only one symbol or letter within pairs. All false-fonts and consonant-strings had five symbols or letters, pseudowords had either four of five letters, and words had either three or four letters. Decoding analysis within stimulus categories were only performed across stimuli with the same number of letters and symbols to prevent length effects. Real word stimuli were selected to have similar log frequency, mean bigram frequency and bigram frequency by position across similar and dissimilar word pairs (measured using the English Lexicon Project^29^). Pseudowords were selected to have similar orthographic neighborhood size and bigram frequency by position across similar and dissimilar pseudoword pairs.

#### Structural MRI acquisition and preprocessing

T1 structural MRIs were used to constrain the cortical source estimates of the current study. Images were acquired with a Siemens 3T Tim Trio system scanner using a magnetization-prepared rapid acquisition with gradient echo sequence (TR = 2100 ms, T1 = 1050 ms, TE = 3.42 ms. 8° flip angle, 256×256×192 acquisition matrices, FOV = 256 mm, and 1 mm isotropic voxels). Cortical surface reconstructions were extracted via Freesurfer^30^.

#### MEG acquisition, preprocessing, and source localization

MEG data were collected on an Elekta Neuromag VectorView MEG system (Elekta Oy, Helsinki, Finland) with 306 sensors (triplets of two orthogonal gradiometers and one magnetometer). Data were sampled at 1000 Hz with simultaneous recording of head position, electrooculogram, and electrocardiogram which were all corrected for off-line. The data were processed with temporal signal-space separation^31^, a 1-50 Hz bandpass filter, and down-sampled to 250 Hz for subsequent analyses.

Minimum norm estimate (MNE) software^32^ was used to project the sensor data onto Freesurfer cortical reconstructions. Regions of interest were manually drawn around the left fusiform gyrus for each subject. Single compartment boundary-element models were calculated from the Freesurfer segmentation and used to compute forward solutions separately for each block, taking shifts in head position into account. Noise covariance matrices were computed from the inter-stimulus interval period, 500 to 30 ms prior to each stimulus presentation. Inverse operators were constructed using the computed noise covariance and forward solutions to obtain source estimates for approximately 7,600 vertices on the cortical surface reconstruction of each subject. Because magnetic sources originating from cortical neurons are typically normal to the cortical surface, tangential source components were scaled by a factor of .4 during the calculation of the inverse solution^33,34^. This procedure resulted in activity of 50-150 lmVTC sources during the category localizer and word-individuation tasks for each subject.

#### Identification of word-sensitive lmVTC sources

Sources in the lmVTC were screened for word selectivity using four-way support vector machines [SVM] applied to 100 ms sliding time windows independently for each source. If the d’ sensitivity index, defined as the inverse of the cumulative normal distribution for true positives for words minus the inverse of the cumulative normal distribution for false positives for words, exceeded chance with p<.05 (uncorrected) a particular source it was considered “word-selective” and belonging to the lmVTC. This yielded a mean +/- standard deviation of 42.4 +/- 32.7 word-selective channels per subject. Only these sources were used for word-individuation decoding. Figure 1 shows the location of these word-sensitive sources across the group. Figure S1 shows the mean event-related field of these word-selective sources to the different stimuli presented in the category localizer task.

**Figure 1.**
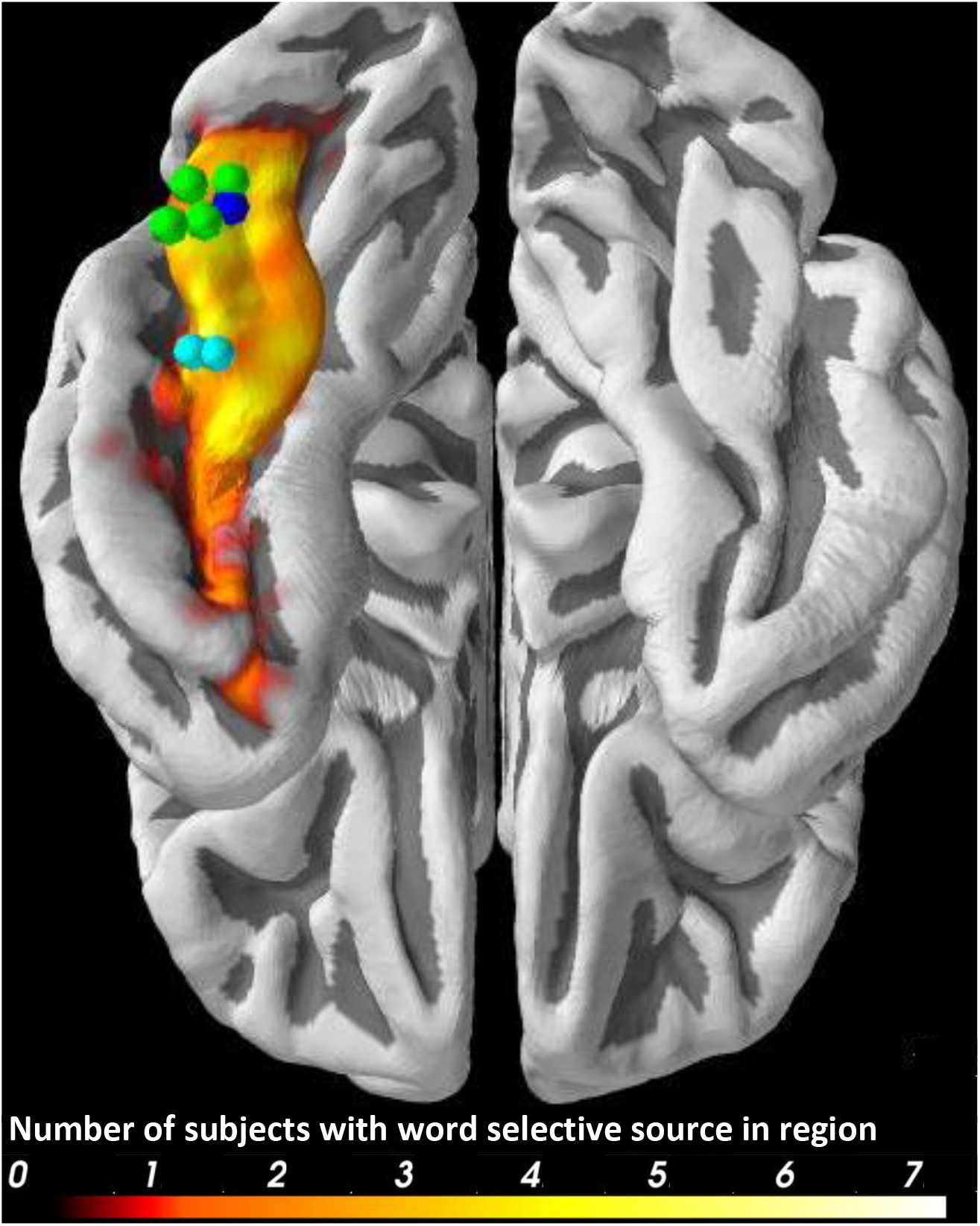
Word selective electrodes and sources in Montreal Neurological Institute common space. Dots are electrodes from the three iEEG patients (P1-green, P2-dark blue, P3-light blue). Number of MEG subjects with word-selective sources localized to a given region of the fusiform represented by color intensity. All sources are constrained to the left fusiform gyrus of the individual’s anatomy.

### Intracranial EEG data collection and preprocessing

#### Participants

Three right-handed patients (2 females, ages 38-64) with intractable epilepsy were included in the study. Inclusion was based on iEEG coverage in left mid-ventral temporal cortex that demonstrated selectivity to words over the other stimulus categories, as defined by the broadband gamma response, event-related potential amplitude, and d’ sensitivity index in an independent category localizer containing words, faces, bodies, houses, hammers, and phase-scrambled objects. Figure S2 shows the d’ sensitivity of each word-selective electrode. Figure S3 shows the event related potential or broadband response of each word-selective electrode to words and other object categories presented during the category localizer task. Figure 2 shows the average d’ sensitivity of the seven word-sensitive electrodes identified across the three subjects. These electrodes were localized using either post-operative T1 structural MRI’s or CT scans. Figure 1 illustrates the location of the word-sensitive electrodes in Montreal Neurological Institute (MNI) stereotaxic-space. Figure S4 illustrates the location of each electrode on the individual patient’s anatomy. None of the electrodes presented here demonstrated ictal activity during the recording sessions, nor were they near the patient’s seizure onset zone. One of the three patients (P1) was included in a previous study^7^; however, data from non-word orthographic stimuli in P1 were not previously reported. The other two participants from that previous study were not shown non-word stimuli, and therefore are not reported here. All patients gave written informed consent under protocols approved by the University of Pittsburgh Medical Center’s Internal Review Board.

**Figure 2.**
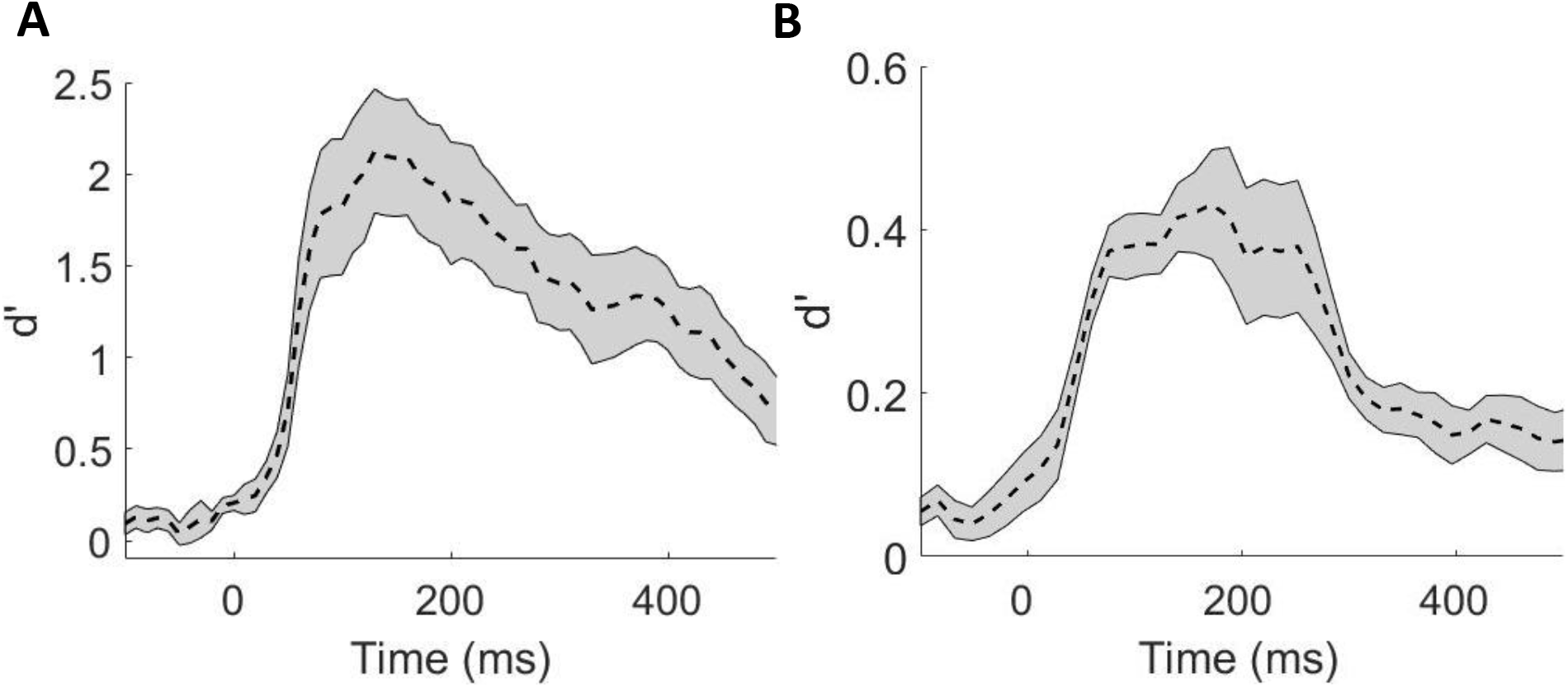
Sensitivity of word-selective electrodes and sources. A) Average sensitivity (*norminv(true positive for words) – norminv(false positive for words*)) of word-selective electrodes in a six-way SVM classifier across all three iEEG participants. Grey represents standard error from the mean across all electrodes. B) Average sensitivity across word-selective MEG sources in lmVTC in a four-way SVM classifier. Grey represents standard error across all subjects.

#### Experimental Paradigm

The experimental paradigm for intracranial subjects was the same as that of the MEG participants besides the following differences: The intracranial category localizer consisted of words, bodies, faces, hammers, houses, and phase scrambled objects. Stimulus on-times for both the category localizer and word-individuation task were increased to 900 ms with 1.8 s mean inter-stimulus interval. The word-individuation task contained the same stimuli as the MEG version; however, to maximize the number of trials per remaining stimuli, consonant-strings were dropped from the stimulus set. Overall, there were approximately 25, 45, and 30 trials per word-individuation stimulus for P1, P2 and P3 respectively--varying according to number of blocks of the task completed.

#### iEEG acquisition and preprocessing

Local field potentials were collected using a Grapevine Neural Interface Processor (Ripple, LLC) at 1000 Hz. Data was bandpass filtered offline from 0.2-115 Hz and notch filtered to exclude 60 Hz line noise using a fourth-order Butterworth filter implemented with FieldTrip^35^. In addition to this, broadband gamma amplitude, defined as the average increase in power from 40-100 Hz, was extracted and normalized to baseline (from 300-50 ms prior to stimulus presentation). Trials with peak amplitudes exceeding 5 standard deviations above or below the mean or exceeding 350 microvolts were eliminated to reduce potential artifacts.

### Multivariate Temporal Pattern Analysis

Data from MEG sources and iEEG electrodes identified as word-selective in the category-localizer task were used for all possible pairwise decoding of the word-individuation stimuli. For example, all word-selective sources in one subject were used as features to a two-class, 3-fold cross-validated SVM classification problem applied to two independent time windows to determine whether the participant was seeing stimulus A or B. Time windows were chosen based on the results from our previous study^7^: 50-200 ms and 200-350 ms for early and late stages of lmVTC processing. LIBLINEAR^36^ was used to implement the SVMs. This resulted in classification accuracy and d’ sensitivity for each pairwise classification problem (44 x 43 / 2) across both time windows. We choose to report d’ sensitivity here because it is normally distributed, unlike classification accuracy, which allows for parametric statistical testing across subjects. Additionally, d’ sensitivity captures effect sizes on the same scale as Cohen’s d, making it easily interpretable. Pairwise d’ sensitivities in the early and late time windows are averaged according the contrast of interest. For example, when determining the classifier sensitivity to real words versus false fonts, all possible pairwise d’ sensitivities between word and false font stimuli were averaged to create one average d’ sensitivity per subject per time window.

Statistical significance of classification accuracy was determined via non-parametric permutation tests. Specifically, category labels were permuted randomly across each pairwise comparison then the two-class SVM was trained on data from the randomly permuted class labels. Classification accuracy for both time windows were computed for 1000 random permutations on the iEEG data and MEG data then averaged over the contrast of interest. Maximum classification accuracy across both time windows was used to construct the null-distribution of classification accuracy and then compared with the corresponding real-label time course. For the MEG data, to obtain the statistical significance of group-wise average classification accuracy, the permuted time courses were also averaged across subject before calculating the maximum accuracy for each of the 1000 random permutations.

### Functional connectivity analysis

Functional connectivity analysis was carried out on the MEG data to evaluate the connectivity dynamics of the lmVTC to the rest of cortex that facilitates real word individuation. Specifically, activity of word-selective sources in lmVTC were averaged and phase-locking values (PLV)^37^ were calculated between this activity and all other cortical sources during the word-individuation task. PLVs were normalized by taking their square root and standardizing relative to a baseline period from 500 to 0 ms prior to stimulus presentation^38^.

Numerous previous non-invasive EEG studies have demonstrated functional connectivity differences in the delta, theta, alpha, and beta frequency bands related to various aspects of reading^39–41^. Therefore, we hypothesized communication between the lmVTC and rest of the language network would be most likely to occur in this frequency range. However, when calculating the phase of low frequency oscillations (i.e. delta and theta bands) using wavelets, these calculations have less temporal resolution than the higher frequency alpha and beta components. Therefore, transformed PLVs were averaged over only the canonical alpha and beta frequency bands (8-30 Hz) then co-registered to the MNI common brain.

To determine spatiotemporal clusters of sources whose PLV to the lmVTC was significantly greater than baseline during the transition from course to fine representations (which occurred at approximately 175-225 ms post-stimulus presentation), cluster statistics were determined via a within-subjects permutation test. Specifically, t-statistics were computed for each source and clustered based on adjacency in time and cortical space. The sum of t-values within each cluster was then compared with the maximum cluster t-value of 10,000 randomly generated sign flipped matrices. This procedure has been shown to effectively correct for multiple spatiotemporal comparisons^42^.

## Results

### Decoding real words from other orthographic stimuli

A support vector machine was trained to discriminate between pairs of word versus other orthographic stimuli using both word sensitive MEG sources and word sensitive iEEG electrodes in lmVTC. Real-words could be discriminated from false fonts, consonant-strings and pseudowords from MEG source activity during both the early (50-200 ms) and late (200-350 ms) time windows (p < .001). Comparable results were seen for word selective iEEG electrodes in the lmVTC besides null results for words versus pseudowords in both windows for patient 3 (Table 1).

**Table 1.** Decoding real words from other stimuli (mean pairwise d’ sensitivity)

|  |  |  |  |  |  |
| --- | --- | --- | --- | --- | --- |
| False fonts vs real words |  |  | Pseudowords vs real words |  |  |
|  | 50-200 ms | 200-350 ms |  | 50-200 ms | 200-350 ms |
| P1 | 0.80*** | 1.5 *** | P1 | 0.40*** | 1.1*** |
| P2 | 0.72 *** | 1.3 *** | P2 | 0.36*** | 0.54 *** |
| P3 | 0.13*** | 0.11** | P3 | 0.046 | -0.034 |
| MEG | 0.26*** | 0.40*** | MEG | 0.051** | 0.16*** |
| Consonant-strings vs real words |  |  |  |  |  |
|  | 50-200 ms | 200-350 ms |  |  |  |
| MEG | 0.17*** | 0.28*** |  |  |  |

\* $p < .05$ \*\* $p < .01$ \*\*\* $p < .001$

A one-way ANOVA indicated a significant difference between mean MEG d’ sensitivity for false fonts, consonant strings, and pseudowords versus real words in the early time window (F = 5.29, p < .01). A post-hoc t-test demonstrated that false fonts versus real words displayed higher d’ sensitivity than pseudowords versus real words across MEG participants in the early time window(p < .01, Bonferroni corrected). Sensitivity for consonant strings versus real words was not significantly different than either false fonts versus real words (p > .2, Bonferroni corrected) or pseudowords versus real words (p > .5) in the early time window. A one-way ANOVA on the late time window also revealed a significant difference between mean MEG d’ sensitivity for false-fonts, consonant strings, and pseudowords versus real words (F = 3.47, p < .05). A post-hoc t-test demonstrated that false fonts versus real words displayed higher d’ sensitivity than pseudowords versus real words across MEG participants during the late time window (p < .05, Bonferroni corrected). However, there was no significant difference between the decoding accuracy of consonant strings versus real words and false-fonts versus real words (p > .5, Bonferroni corrected) or pseudowords versus real words (p > .6, Bonferroni corrected) in the late time window.

### Decoding orthographically similar and dissimilar stimuli

First we determined if we could replicate our previous iEEG results regarding the dynamics of similar and dissimilar individual word decoding^7^ using source localized MEG. Using activity evoked from word-selective MEG sources in lmVTC, orthographically dissimilar real words could be decoded from one another in the early and late time windows (p < 0.001 in both windows, see Table 2). However, orthographically similar real words could not be significantly decoded from each other until the late time window from 200-350 ms (p < .05). A similar pattern was seen in 4 of the 5 iEEG subjects (2/3 reported here and 3/3 reported previously^7^, with one subject shared between the two studies, see Methods). This pattern is consistent with our previous iEEG results regarding the dynamics of similar and dissimilar individual word decoding^7^.

**Table 2.** Decoding orthographically similar and dissimilar stimuli (mean pairwise d’ sensitivity)

|  |  |  |  |  |  |  |
| --- | --- | --- | --- | --- | --- | --- |
| Orthographically similar real words |  |  | Orthographically dissimilar real words |  |  |  |
|  | 50-200 ms | 200-350 ms |  | 50-200 ms | 200-350 ms |  |
| P1 | -0.0006 | 0.56** | P1 | 0.41*** | 0.96*** |  |
| P2 | -0.075 | 0.85*** | P2 | 0.62*** | 0.60*** |  |
| P3 | -0.13 | -0.26 | P3 | -0.12 | -0.028 |  |
| MEG | -0.048 | 0.10* | MEG | 0.091*** | 0.14*** |  |
| Orthographically similar pseudowords |  |  | Orthographically dissimilar pseudowords |  |  |  |
|  | 50-200 ms | 200-350 ms |  | 50-200 ms | 200-350 ms |  |
| P1 | -0.049 | 0.040 | P1 | 0.29** | 0.72*** |  |
| P2 | -0.041 | 0.089 | P2 | 0.17 | 0.19* |  |
| P3 | -0.0002 | 0.061 | P3 | 0.48054 | 0.016 |  |
| MEG | 0.087 | 0.040 | MEG | 0.090** | 0.13*** |  |
| Orthographically similar consonant-strings |  |  | Orthographically dissimilar consonant-strings |  |  |  |
|  | 50-200 ms | 200-350 ms |  | 50-200 ms | 200-350 ms |  |
| MEG ONLY | -0.079 | 0.048 | MEG ONLY | 0.040 | 0.13*** |  |
| Orthographically similar false fonts |  |  | Orthographically dissimilar false fonts |  |  |  |
|  | 50-200 ms | 200-350 ms |  | 50-200 ms | 200-350 ms |  |
| P1 | 0.029 | 0.43* | P1 | 0.13* | 0.70*** |  |
| P2 | 0.47* | -0.092 | P2 | 0.11 | 0.35*** |  |
| P3 | -0.049 | -0.083 | P3 | -0.018 | 0.19* |  |
| MEG | 0.017 | 0.069 | MEG | 0.10*** | 0.18*** |  |

\* $p < .05$ \*\* $p < .01$ \*\*\* $p < .001$

Like real words, orthographically dissimilar pseudowords (p<.01 early, p<.001 late), consonant-strings (p<.001 only in the late time window), and false fonts (p<.001 in both windows) could be significantly discriminated within lmVTC activity using MEG and in most iEEG cases, particularly in the late time window (Table 2). In contrast to real words, orthographically similar consonant-strings, pseudowords and false fonts could not be consistently decoded with MEG or iEEG at either time window (Table 2).

### Functional connectivity between the lmVTC and rest of the brain during the disambiguation of real word stimuli

An open question is whether the individuation of real word stimuli is achieved locally within lmVTC or involves a network level process that lmVTC contributes to. To determine functional interactions that occur during the transition from coarse to fine orthographic representations in the lmVTC, phase locking values (PLVs) were calculated between the average activity of word-selective sources localized to lmVTC and the rest of the cortical sources from 175-225 ms post stimulus presentation. Normalizing these PLVs with respect to baseline and averaging over canonical alpha and beta frequency bands (see Methods) gave two clusters of sources that demonstrated above chance connectivity relative to pre-stimulus baseline at the cluster-level (p<.05). Figure 3 illustrates the spatial locus of these clusters, which includes one that extends from right early visual cortex to right lateral occipital cortex and one encompassing a region anterior to the left fusiform gyrus, extending from the parahippocampal gyrus to the inferior temporal sulcus. No significant clusters were seen when directly contrasting the PLVs evoked by real words to those evoked by pseudowords, consonant strings, or false-fonts.

**Figure 3.**
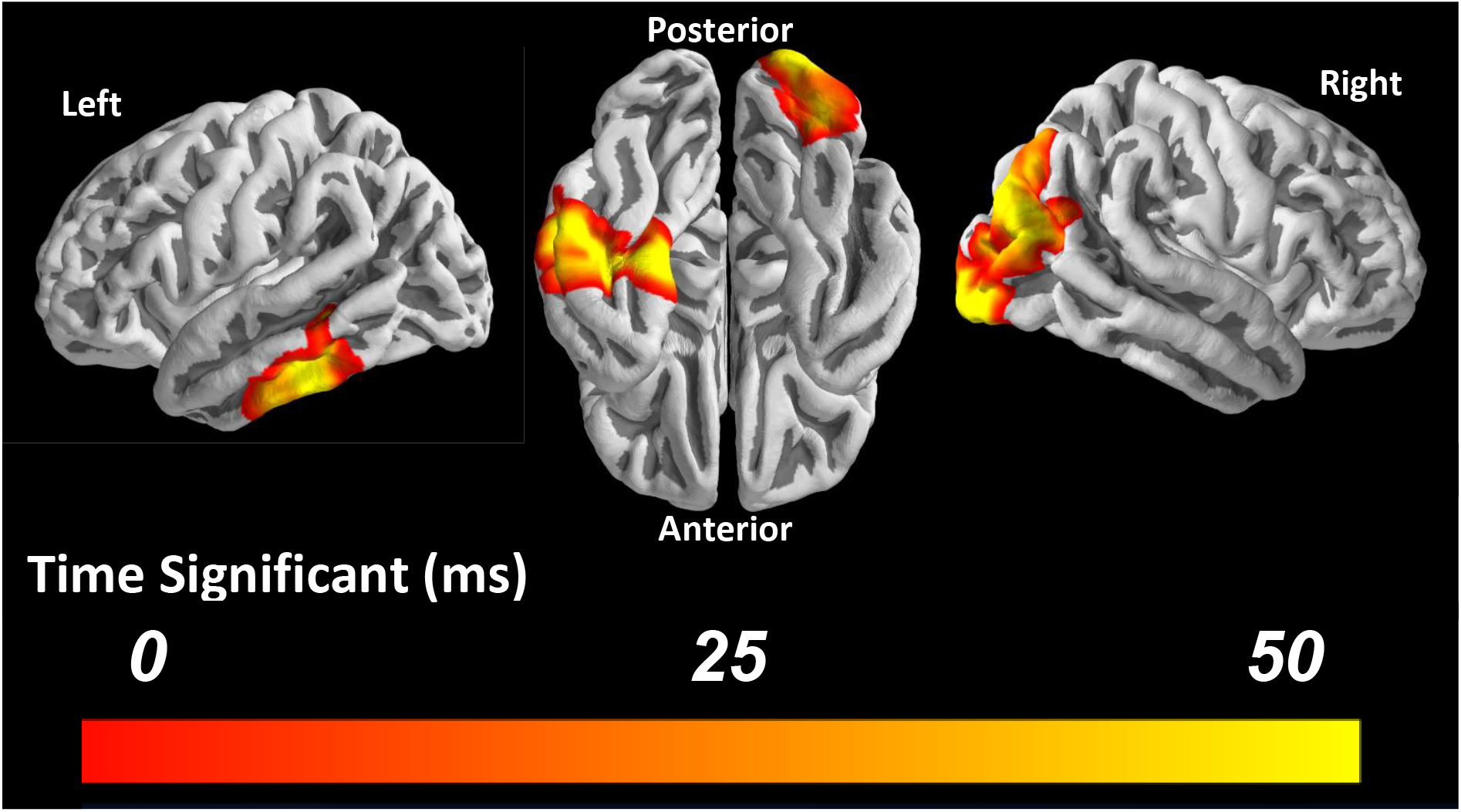
Spatiotemporal clusters of significant phase-locking values (PLV) to the word-selective sources in lmVTC during the individuation of real-word representations (175-225 ms). Color intensity illustrates the duration each source had elevated PLV (p<.01, uncorrected) during real word trials versus baseline with 50 ms being the maximum possible duration. Significant clusters include one in right early visual cortex and in the left anterior temporal lobe.

## Discussion

We found that lmVTC representations of real words, pseudowords, consonant strings and false fonts were all initially coarse, allowing only for decoding of visually dissimilar stimuli from each other. Real word representations in the lmVTC became disambiguated over time, allowing for reliable decoding of orthographically similar real words from lmVTC activity after 200 ms post-stimulus presentation. However, decoding of visually similar pseudowords, letter strings and false fonts from lmVTC activity did not rise above chance during either the early or late time window. Finally, in the transition period between coarse and individuated representations of real words, we observed significant functional connectivity between the lmVTC and early visual cortex and between the lmVTC and more anterior regions of the left temporal lobe.

Our decoding analyses for orthographically similar versus dissimilar real words replicate a previous finding from our group: real word representations in the lmVTC are initially coarse but disambiguate with time to allow for the reliable representation of orthographically similar real words^7^. This replication, in addition to the high correspondence between the MEG and iEEG results presented here, provide an important cross-validation of iEEG and source-localized MEG. Specifically, these results demonstrate the sensitivity of MEG to subtle stimulus-induced changes in neural activity and source-localization’s ability to approximately identify the neuroanatomical origins of those neural signatures. Furthermore, the correspondence between source-localized MEG and iEEG validate iEEG results using data from a healthy population with larger sample size. Additionally, MEG supplements sensitive iEEG data with full brain coverage, facilitating analyses that require broader coverage, like the functional connectivity analysis presented here. Thus, combining iEEG and MEG with similar experiments is a potentially powerful paradigm to cross-validate, replicate, and extend findings by leveraging the respective strengths of these two recording techniques.

By combining MEG and iEEG in the current study, we were able to demonstrate that early, coarse coding in the lmVTC exists not only for real words, but also false fonts, consonant-strings and pseudowords. This early representation may support rapid disambiguation of orthographically dissimilar stimuli. However, our data suggests that these coarse lmVTC representations are subsequently disambiguated through interactions between visual and semantic networks, which allows the reliable individuation of orthographically similar real words. Given that we were only able to decode visually similar real words from late lmVTC activity, and not visually similar stimuli from other orthographic categories, this suggests that lmVTC representations of learned word forms are individuated to a greater degree than unfamiliar orthographic entities.

This conclusion is supported by previous studies observing decreases in lmVTC BOLD responses for learned relative to unfamiliar orthographic entities^16,19^, which may reflect more individuated lmVTC representations for learned word forms in the later stages of processing. Further, the time-course of real word individuation in the lmVTC is supported by scalp EEG evidence demonstrating lexical-semantic influences on visual word recognition that are observed approximately 250 ms post-stimulus presentation, as shown by studies involving transposed letter^43^ and morphological primes^44^. However, the current study cannot rule out the possibility that the lmVTC has the capacity to individuate unfamiliar orthographically stimuli, either through purely bottom-up visual mechanisms or interactions with phonological processing networks, since care must be taken when interpreting null results. With that in mind, the results reported here do suggest that these stimuli are likely represented less robustly than known word forms.

Our results, which support early, coarse orthographic coding in the lmVTC, contrasts with previous results obtained from rapid adaptation functional magnetic resonance imaging (fMRI) studies. Glezer et al. 2009^18^ reported no effect of orthographic similarity on the on the adaptation of BOLD response to successively presented real words. Based on these results, the authors suggested that lmVTC representations of real words are not coarse, but rather based on individuated whole word templates, hierarchically assembled from rapid, bottom-up visual information processing^18^. The results reported here show that the early response of lmVTC is coarse, potentially reflecting an orthographic similarity space^45^. It has been suggested that the early response of an area reflects its intrinsic coding, since later activity is more susceptible to top-down and network-level influences^3^. Thus, these results suggest that lmVTC’s intrinsic code does not reflect individuated whole word templates. A potential explanation for the conflicting results is the difference in temporal resolution afforded by fMRI relative to MEG and iEEG. The sluggish hemodynamic response measured by fMRI may be disproportionately sensitive to the late stage of lmVTC processing, when visually similar real words can be disambiguated from each other. Our results suggest that the early representations in the lmVTC, which potentially arise from bottom-up visual processing, are consistent with coarse orthographic coding. Whole word representations emerge in the lmVTC over time, however they likely require network interactions with semantic and visual regions to reliably disambiguate orthographically similar real words^25^.

Functional connectivity analyses presented also support this conclusion. During real word trials, differences in phase-locking between two spatially distinct clusters, one centered on early visual areas (early visual cortex and right lateral occipital cortex) and the other on the left anterior temporal lobe, were present during the 50 ms transition from early to late decoding windows. This suggests that the individuation of real word representations in the lmVTC takes place through recurrent interactions between regions both earlier in the ventral visual hierarchy and higher-level processing regions. Interactions with occipital regions may reflect continued accumulation of visual information over time, while anterior temporal regions may contribute learned information about real words that support the disambiguation of word forms through semantic properties which are largely orthogonal to the orthographic properties of printed words^25^. This role for the anterior temporal cortex in reading is supported by studies demonstrating increased BOLD activation of this regions to pseudowords trained to have semantic associations^46^ and studies of sentence comprehension^47^.

However, no clusters of lmVTC functional connectivity were found to be significantly different between real words and the other word-like stimuli at the group level. Thus, it may be that neural communication is shared among a similar set of regions regardless of the stimuli, but only supports individuated representations in the lmVTC if there is useful stored information in a given node of the network. Notably, these functional connectivity results suggest that individuated representations are an emergent property of network interactions, with multiple nodes of the network contributing to and reflecting individuated representations. Thus, individuation of real words in the lmVTC is unlikely to be a result of solely visual processing occurring in this region, but rather a local reflection of a network-level computation. A similar timing pattern has been reported for face individuation in the fusiform gyrus, where early activity in response to faces is coarse and later activity is individuated^48^ and may also reflect network-level interactions^49^. This suggests that a similar dynamic process is conserved between both word and face stimuli and may reflect a general principle of visual processing for other visual stimuli as well.

Taken together, our results support the idea of an early, coarse code in the lmVTC that is sharpened through recurrent interactions between occipital and anterior temporal regions. First, a coarse level representation in the lmVTC, built through bottom-up visual processing, allows for decoding of visually dissimilar stimuli within 200 ms of stimulus presentation. Next, interactions between anterior ventral temporal regions, possibly containing stored knowledge about words, and low-order visual regions assist in disambiguating real word representations over time. This information ultimately allows the individuation of visually similar real words. Further work investigating to what degree this process is sensitive to word context (i.e. when a word is presented in a meaningful sentence) and whether individuated word representations are conveyed throughout the language network is necessary to better understand the computations which facilitate expert reading.

## Acknowledgements

We would like to thank the patients and staff at the epilepsy monitoring unit at the University of Pittsburgh Medical Center, without whom this research would not be possible. This work was supported by National Institute of Health Award NIH T32NS007433-20 (to M.J.B.), National Institute of Mental Health Award NIH R01MH107797 (to A.S.G), National Science Foundation Award 1734907 (to A.S.G.), and Eunice Kennedy Shriver National Institute of Child Health and Human Development Award R01HD060388 (to J.A.F.).

## Author Contributions

Conceptualization, M.J.B., Y.L., J.A.F., A.S.G., and E.A.H.; Methodology, M.J.B., J.A.F., A.S.G., and E.A.H.; Investigation, M.J.B., E.A.H., Y.L., M.R.R., and M.J.W.; Formal Analysis, M.J.B., E.A.H., and Y.L.; Writing – Original Draft, M.J.B. and A.S.G.; Writing – Review & Editing, M.J.B., E.A.H., J.A.F., M.R.R., and A.S.G.; Resources, M.R.R. and A.S.G.; Funding Acquisition, M.J.B., J.A.F., and A.S.G.

## References

1. Cohen, L. et al. Language‐specific tuning of visual cortex? Functional properties of the Visual Word Form Area. Brain 125, 1054–1069 (2002).

2. McCandliss, B. D., Cohen, L. & Dehaene, S. The visual word form area: expertise for reading in the fusiform gyrus. Trends Cogn. Sci. 7, 293–299 (2003).

3. Dehaene, S. & Cohen, L. The unique role of the visual word form area in reading. Trends in Cognitive Sciences 15, 254–262 (2011).

4. Gaillard, R. et al. Direct Intracranial, fMRI, and Lesion Evidence for the Causal Role of Left Inferotemporal Cortex in Reading. Neuron 50, 191–204 (2006).

5. Price, C. J. & Devlin, J. T. The Interactive Account of ventral occipitotemporal contributions to reading. Trends in Cognitive Sciences 15, 246–253 (2011).

6. Devlin, J. T., Jamison, H. L., Gonnerman, L. M. & Matthews, P. M. The role of the posterior fusiform gyrus in reading. J. Cogn. Neurosci. 18, 911–22 (2006).

7. Hirshorn, E. A. et al. Decoding and disrupting left midfusiform gyrus activity during word reading. Proc. Natl. Acad. Sci. 113, 8162–8167 (2016).

8. Pflugshaupt, T. et al. About the role of visual field defects in pure alexia. Brain 132, 1907–1917 (2009).

9. Leff, A. P. et al. The functional anatomy of single-word reading in patients with hemianopic and pure alexia. Brain 124, 510–21 (2001).

10. Mani, J. et al. Evidence for a basal temporal visual language center: cortical stimulation producing pure alexia. Neurology 71, 1621–7 (2008).

11. Dehaene, S. et al. How learning to read changes the cortical networks for vision and language. Science 330, 1359–64 (2010).

12. Shaywitz, B. A. et al. Development of left occipitotemporal systems for skilled reading in children after a phonologically- based intervention. Biol. Psychiatry 55, 926–933 (2004).

13. Schlaggar, B. L. & McCandliss, B. D. Development of Neural Systems for Reading. Annu. Rev. Neurosci. 30, 475–503 (2007).

14. Brem, S. et al. Brain sensitivity to print emerges when children learn letter-speech sound correspondences. Proc. Natl. Acad. Sci. U. S. A. 107, 7939–44 (2010).

15. Ben-Shachar, M., Dougherty, R. F., Deutsch, G. K. & Wandell, B. A. The Development of Cortical Sensitivity to Visual Word Forms. J. Cogn. Neurosci. 23, 2387–2399 (2011).

16. Glezer, L. S., Kim, J., Rule, J., Jiang, X. & Riesenhuber, M. Adding words to the brain’s visual dictionary: novel word learning selectively sharpens orthographic representations in the VWFA. J. Neurosci. 35, 4965–72 (2015).

17. Stevens, W. D., Kravitz, D. J., Peng, C. S., Tessler, M. H. & Martin, A. Privileged Functional Connectivity between the Visual Word Form Area and the Language System. J. Neurosci. 37, 5288–5297 (2017).

18. Glezer, L. S., Jiang, X. & Riesenhuber, M. Evidence for Highly Selective Neuronal Tuning to Whole Words in the “Visual Word Form Area”. Neuron 62, 199–204 (2009).

19. Kronbichler, M. et al. The visual word form area and the frequency with which words are encountered: evidence from a parametric fMRI study. Neuroimage 21, 946–953 (2004).

20. Binder, J. R., Medler, D. A., Westbury, C. F., Liebenthal, E. & Buchanan, L. Tuning of the human left fusiform gyrus to sublexical orthographic structure. Neuroimage 33, 739–748 (2006).

21. Dehaene, S. C., Le Clec’H, G., Poline, J.-B., Le Bihan, D. & Cohen, L. The visual word form area: a prelexical representation of visual words in the fusiform gyrus. Neuroreport 13, 321–325 (2002).

22. Annelies Baeck, Dwight Kravitz, Chris Baker & Hans P. Op de Beecka. Influence of lexical status and orthographic similarity on the multi-voxel response of the visual word form area. Neuroimage 111, 321–328 (2015).

23. Cohen, L. et al. The visual word form area: spatial and temporal characterization of an initial stage of reading in normal subjects and posterior split-brain patients. Brain 123 (Pt 2), 291–307 (2000).

24. Price, C. J. & Devlin, J. T. The myth of the visual word form area. Neuroimage 19, 473–481 (2003).

25. Plaut, D. C., Mcclelland, J. L., Seidenberg, M. S., Patterson, K. & Program, N. Understanding Normal and Impaired Word Reading: Computational Principles in Quasi-Regular Domains. Psychological Review (1996).

26. Fiez, J. A. & Petersen, S. E. Neuroimaging studies of word reading. Proc. Natl. Acad. Sci. U. S. A. 95, 914–21 (1998).

27. Price, C. J. & McCrory, E. Functional Brain Imaging Studies of Skilled Reading and Developmental Dyslexia. in The Science of Reading Handbook (eds. Snowling, M. J. & Hulme, C.) 473–496 (Blackwell Publishing, 2005).

28. Brainard, D. H. The Psychophysics Toolbox. Spat. Vis. 10, 433–436 (1997).

29. Balota, D. A. et al. The English Lexicon Project. Behavior Research Methods 39, (2007).

30. Dale, A. M., Fischl, B. & Sereno, M. I. Cortical Surface-Based Analysis. Neuroimage 9, 179–194 (1999).

31. Taulu, S. & Hari, R. Removal of magnetoencephalographic artifacts with temporal signal-space separation: Demonstration with single-trial auditory-evoked responses. Hum. Brain Mapp. 30, 1524–1534 (2009).

32. Gramfort, A. et al. MNE software for processing MEG and EEG data. Neuroimage 86, 446–460 (2014).

33. Lin, F.-H., Belliveau, J. W., Dale, A. M. & Hämäläinen, M. S. Distributed current estimates using cortical orientation constraints. Hum. Brain Mapp. 27, 1–13 (2006).

34. Lin, F.-H. et al. Assessing and improving the spatial accuracy in MEG source localization by depth-weighted minimum-norm estimates. (2006). doi:10.1016/j.neuroimage.2005.11.054

35. Oostenveld, R., Fries, P., Maris, E. & Schoffelen, J.-M. FieldTrip: Open source software for advanced analysis of MEG, EEG, and invasive electrophysiological data. Comput. Intell. Neurosci. 2011, 156869 (2011).

36. Fan, R.-E., Chang, K.-W., Hsieh, C.-J., Wang, X.-R. & Lin, C.-J. LIBLINEAR: A Library for Large Linear Classification. J. Mach. Learn. Res. 9, 1871–1874 (2008).

37. Lachaux, J.-P., Rodriguez, E., Martinerie, J. & Varela, F. J. Measuring Phase Synchrony in Brain Signals. Hum Brain Mapp. 8, 194–208 (1999).

38. Ghuman, A. S., McDaniel, J. R. & Martin, A. A wavelet-based method for measuring the oscillatory dynamics of resting-state functional connectivity in MEG. Neuroimage 56, 69–77 (2011).

39. Weiss, S. & Mueller, H. M. The contribution of EEG coherence to the investigation of language. Brain Lang. 85, 325–343 (2003).

40. Spironelli, C. & Angrilli, A. Developmental aspects of language lateralization in delta, theta, alpha and beta EEG bands. Biol. Psychol. 85, 258–267 (2010).

41. Weiss, S. & Mueller, H. M. “Too Many betas do not Spoil the Broth”: The Role of Beta Brain Oscillations in Language Processing. Front. Psychol. 3, 201 (2012).

42. Maris, E. & Oostenveld, R. Nonparametric statistical testing of EEG- and MEG-data. J. Neurosci. Methods 164, 177–190 (2007).

43. Duñabeitia, J. A., Molinaro, N., Laka, I., Estévez, A. & Carreiras, M. N250 effects for letter transpositions depend on lexicality: ‘casual’ or ‘causal’? Neuroreport 20, 381–387 (2009).

44. Morris, J., Frank, T., Grainger, J. & Holcomb, P. J. Semantic transparency and masked morphological priming: An ERP investigation. (2007). doi:10.1111/j.1469-8986.2007.00538.x

45. Baeck, A., Kravitz, D., Baker, C. & Op de Beeck, H. P. Influence of lexical status and orthographic similarity on the multi-voxel response of the visual word form area. Neuroimage 111, 321–328 (2015).

46. Malone, P. S., Glezer, L. S., Kim, J., Jiang, X. & Riesenhuber, M. Multivariate Pattern Analysis Reveals Category-Related Organization of Semantic Representations in Anterior Temporal Cortex. J. Neurosci. 36, 10089–96 (2016).

47. Vandenberghe, R., Nobre, A. C. & Price, C. J. The response of left temporal cortex to sentences.pdf. 550–560 (2002).

48. Ghuman, A. S. et al. Dynamic encoding of face information in the human fusiform gyrus. Nat. Commun. 5, 5672 (2014).

49. Li, Y., Richardson, R. M. & Ghuman, A. S. Multi-Connection Pattern Analysis: Decoding the representational content of neural communication. Neuroimage 162, 32–44 (2017).

